## Supplemental Figs S1-S2 and Table S1 for "“Superiority of Rhythmic Auditory Signals over Electrical Stimulation to Entrain Behavior”"

**This PDF file includes:**

Figs S1 to S2
Table S1

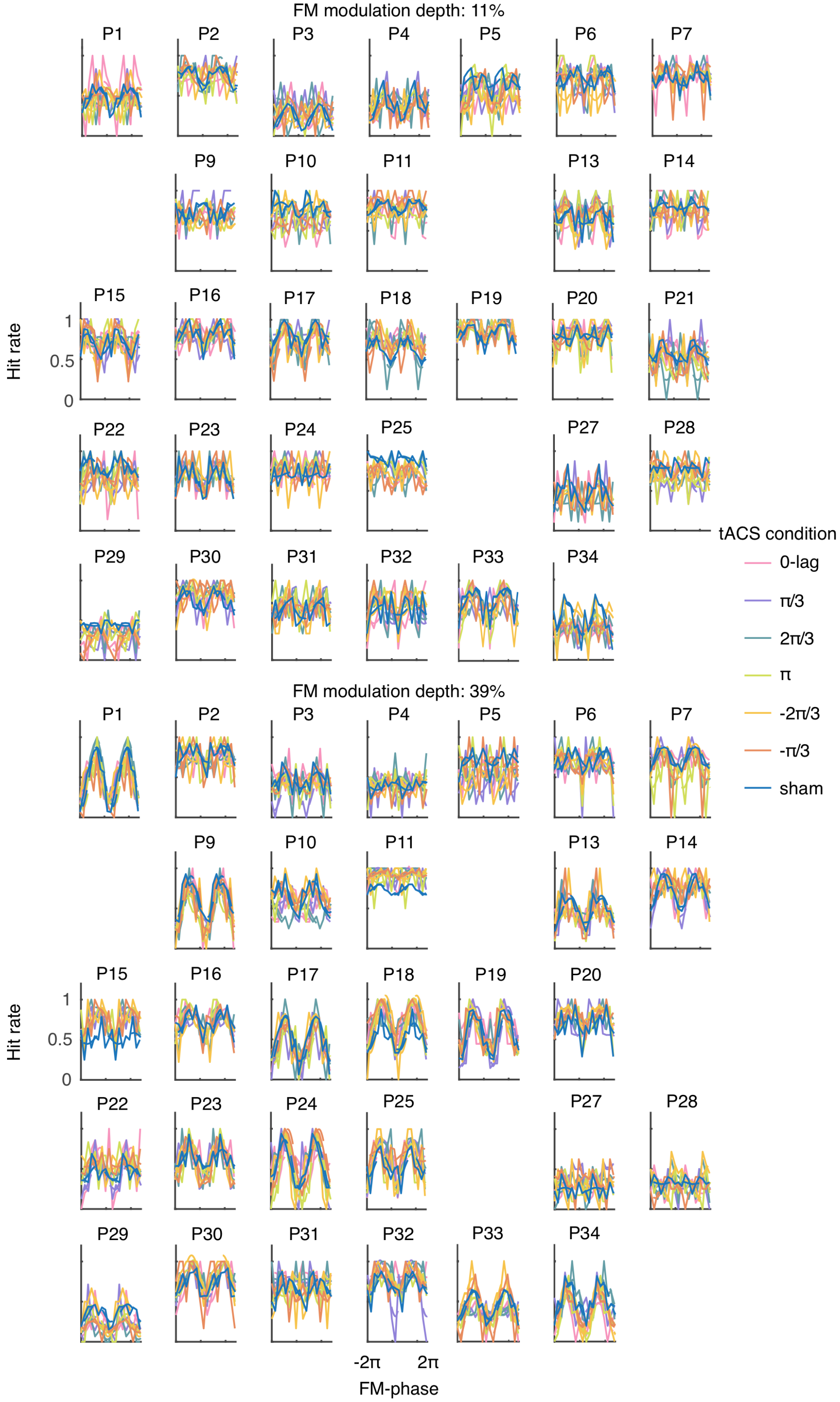


**Fig. S1. Experiment 1.** Supplementary figure accompanying Fig. 1d. Individual data showing hit rates as a function of FM-stimulus phase for each tACS condition when the FM stimulus is modulated at 11% (top) and 39% (bottom). Each plot shows data from a different participant and modulation depth. Empty plots represent participants excluded due to incomplete data.

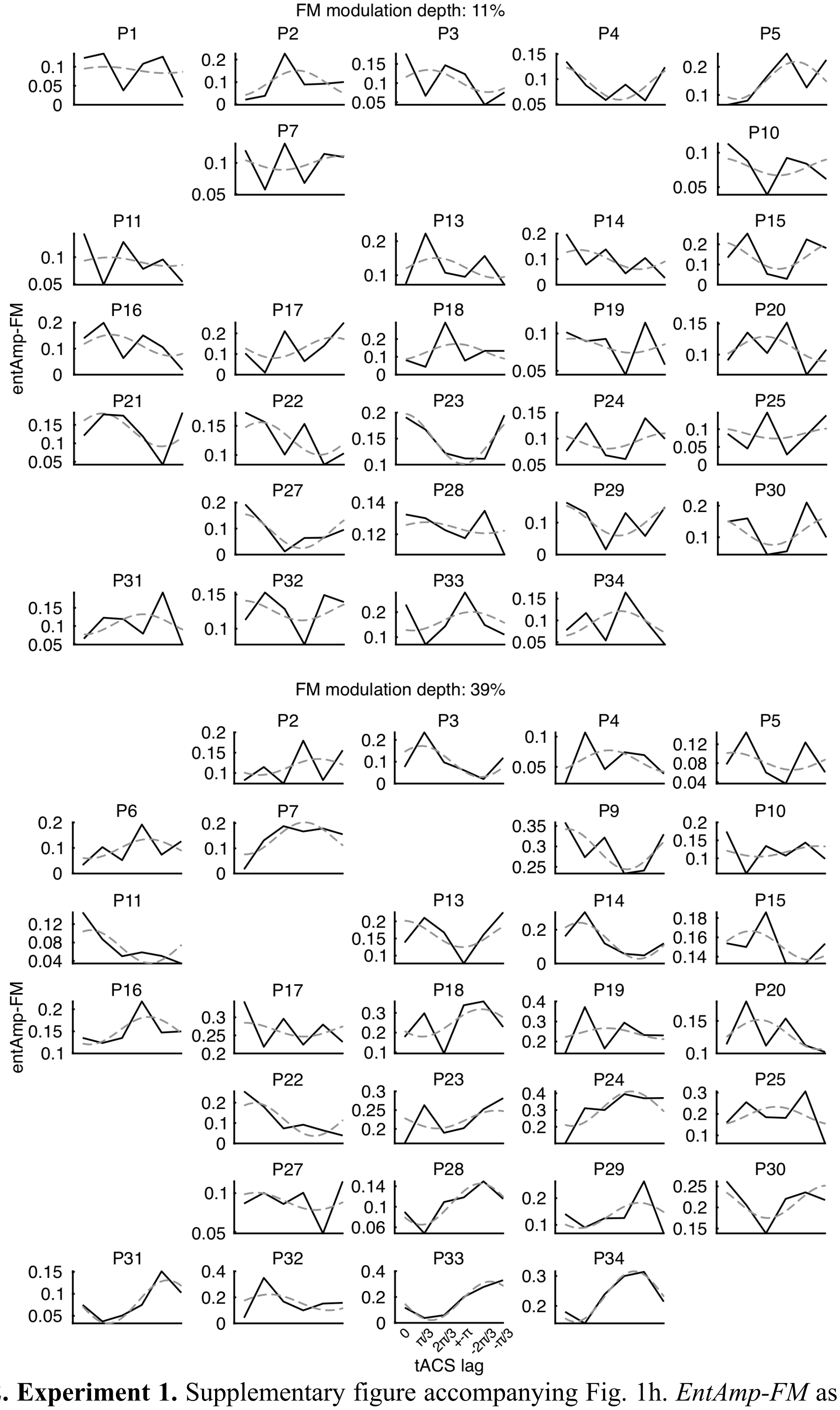


**Fig. S2. Experiment 1.** Supplementary figure accompanying Fig. 1h. *EntAmp-FM* as a function of tACS lag for each modulation depth; 11% (top) and 39% (bottom). Solid lines show the actual amplitude parameters obtained from the initial cosine fits on the data in Fig. 1d and dashed lines represent the second cosine fit to estimate the optimal tACS phase for modulating entrainment to the auditory stimulus. Each plot is a different participant and modulation depth. Empty plots represent participants excluded due to incomplete data.

**Table S1. Mixed effects logistic regression models explaining single trial gap detection performance in Experiment 1**

| | Formula | AIC | Δ AIC* |
|---|---|---|---|
| **1** | ***response ~ 1 + sinAUD + cosAUD + (1 \| subjectNr)*** | 33917.95 | 0 |
| 2 | *response ~ 1 + sintACS + costACS + sinAUD + cosAUD + (1 \| subjectNr) + (1 + sintACS \| subjectNr) + (1 + costACS \| subjectNr)* | 33922.61 | 4.66 |
| 3 | *response ~ 1 + sinAUD*modDepth + cosAUD*modDepth + (1 \| subjectNr)* | 33923.95 | 6.00 |
| 4 | *response ~ 1 + sinAUD + cosAUD + sintACS*modDepth + costACS*modDepth + (1 \| subjectNr) + (1 + sintACS \| subjectNr) + (1 + costACS \| subjectNr)* | 33928.61 | 10.66 |
| 5 | *response ~ 1 + sintACS + costACS + (1 \| subjectNr) + (1 + sintACS \| subjectNr) + (1 + costACS \| subjectNr)* | 34787.58 | 869.63 |

Models are organized from smallest to highest AIC. Δ AIC relative to winning model. The winning model is also highlighted in bold. AIC, Akaike information criterion
